# Quantitative and large-scale investigation of human TCR-HLA Cross-Reactivity

**DOI:** 10.1101/2025.02.20.639349

**Authors:** Mingyao Pan, Yuhao Tan, Yizhou Tracy Wan, Jing Hu, Julia Fleming, Hailong Hu, Ziqi Yang, Xiaowei Zhan, Bo Li

**Affiliations:** Department of Bioengineering, School of Engineering, University of Pennsylvania, Philadelphia, PA, USA; Center for Computational and Genomic Medicine, The Children’s Hospital of Philadelphia, Philadelphia, PA, USA; Department of Pathology and Laboratory Medicine, University of Pennsylvania, Philadelphia, PA, USA; Quantitative Biomedical Research Center, Peter O’Donnell School of Public Health, University of Texas Southwestern Medical Center, Dallas, TX, USA

## Abstract

The interaction between Human leukocyte antigens (HLA) and T cell receptor (TCR) is essential for adaptive immune recognition. While it is known that one TCR can map to multiple HLA alleles, the extent of this cross-reactivity remains poorly understood. Here, we introduce THNet, a TCR-based HLA similarity inference method, and performed a comprehensive analysis of HLA-TCR cross-reactivity. This method is built upon clustering over 47 million TCRs to identify over 9 million significant HLA-TCR pairs. We created similarity networks for both class I and class II HLA alleles, illustrating how peptide cross-presentation contributes to HLA-TCR cross-reactivity. This analysis revealed novel disease-susceptibilities missed by single-HLA enrichment analyses, especially in the Black populations. Finally, we demonstrated that THNet prioritized optimal HLA mismatch candidates for organ transplantation, thereby improving patient survival rates. Our investigation of HLA-TCR cross-reactive network might provide useful insights for autoimmune risk prediction and better transplantation outcomes.

**One Sentence Summary:** We introduced THNet, a large-scale TCR-based HLA similarity mapping network that uncovers previously unrecognized cross-reactivity patterns across HLA alleles and provides valuable insights into their influence on disease susceptibility and graft rejection.

## INTRODUCTION

T lymphocytes are essential components of adaptive immunity, playing critical roles in targeting and eliminating infectious pathogens and tumor cells(*1–3*). The T cell receptors (TCRs) on the surface of T cells enable them to recognize and respond to specific epitopes presented by HLA molecules(*4, 5*). The specificity of TCR recognition, however, is not strictly one-to-one; rather, it follows a many-to-many pattern, where a single TCR can recognize multiple peptide-HLA (pHLA) complexes, and, conversely, a single pHLA complex can activate different T cell clones(*6–11*). This ability of a single TCR to recognize diverse pHLA is termed TCR cross-reactivity, ensuring each individual’s TCR repertoire to recognize the broad array of potential pathogens(*11–13*). Some previous studies suggest that a single TCR on average can bind to over 10⁶ different pHLA complexes(*11, 13*).

TCR cross-reactivity extends beyond individual HLA subtypes, with instances of single T cell clones being activated by allogeneic HLA molecules documented decades ago(*14–19*). Alloreactive T cells, which react to allogeneic HLA molecules, are key mediators of graft rejection(*20–22*), and studies have shown that approximately 45% of viral-specific memory CD4^+^ and CD8^+^ T cells can cross-react with at least one allogeneic HLA molecule(*23–25*). This TCR-HLA cross-reactivity has been implicated as the cause of selected autoimmune disorders(*23, 26–29*). For instance, the cross-reactivity of human T cell clone Hy.2E11 between Epstein-Barr virus (EBV)-derived peptide presented by DRB5*01:01 and human myelin basic protein presented by DRB1*15:01 underlies the association between EBV infection and an increased risk of multiple sclerosis(*26, 27, 30*). Additional examples of TCRs with identical alpha and beta chains recognizing peptides presented by different HLA alleles can also be found in various TCR antigen-specificity databases, such as IEDB and VDJdb(*31, 32*). Due to the experimental challenges of pairing TCRs with their cognate antigens, the full extent of cross-reactivity across different HLA alleles remains poorly understood, despite decades of research highlighting its significance in various disease settings. A more comprehensive and systematic investigation is therefore needed to better understand the degree and mechanisms of TCR-HLA cross-reactivity.

An individual’s HLA composition significantly influences TCR clonotype diversity and clonal expansion(*33–35*). During thymic selection, different HLA alleles select distinct subsets of thymocytes to develop into functional T cells(*36–38*). Beyond thymus, different HLAs present distinct peptides derived from external pathogens or cancer cells and activate antigen-specific T cell populations(*39–41*). As a result, HLA alleles are among the most critical genes that regulate T cell responses and shape TCR repertoire(*41–43*), with some alleles exhibiting strong disease associations(*35, 42, 44–47*). Many efforts have been devoted to studying HLA-TCR associations(*48–51*), but the complexity of the human HLA gene cluster and remarkable HLA polymorphism present significant challenges in understanding the effects of individual HLA alleles on TCR repertoire. At the individual level, each person carries up to six classical HLA class I alleles (HLA-A/B/C) and multiple HLA class II alleles (HLA-DP/DQ/DR)(*52, 53*). At the population level, HLA genes exhibit extensive diversity, with over 28,000 HLA class I and 12,000 HLA class II alleles currently documented in the IPD-IMGT/HLA database(*54*). While most studies regard every HLA allele as a factor independent from each other(*55–58*), some HLA molecules exert overlapping effects on TCR repertoire(*59–62*). Developing a more sophisticated model that accounts for the functional similarities of different HLA alleles could unlock new insights into many HLA-related clinical settings, including disease associations and HLA compatibility in organ transplantation.

Investigations of sequence-based similarity between HLA alleles revealed that the variations in different regions of HLA molecules have significantly different impacts on their functions (*63–65*). Hence, more efforts were focused on the peptide/TCR-contacting regions, to define similar HLA alleles by shared sequences in these functional sites(*62, 66–69*). Nevertheless, even within these sites, the impacts of single amino acid (AA) variations on peptide binding cannot be precisely predicted(*70–72*). Therefore, investigation of HLA sequence similarity cannot adequately represent their peptide-presenting or TCR-activating capabilities. To bridge this gap, rather than analyzing HLA itself, we directly examined their functional consequences, *i.e.* the fraction of TCR repertoire associated with each allele. Specifically, we developed a TCR-based HLA similarity mapping network, THNet, to quantify HLA-TCR cross-reactivity and functional similarity between different HLA alleles. This is achieved through ultra-fast clustering(*73*) of over 47 million TCRs from individuals with known HLA genotypes. This method allowed us to group HLA alleles into functionally related networks and to identify novel HLA disease susceptibilities in the minority populations. Moreover, THNet assisted in selecting optimal HLA mismatch candidates within limited donor pools to minimize the risk of anti-graft T cell responses, serving as a new prognostic modality for organ transplantation. In summary, our systematic and quantitative investigation of TCR-based HLA similarity provided valuable insights into how different HLA alleles influence outcomes in various clinical settings.

## RESULTS

### The THNet framework

To quantitatively investigate TCR-HLA cross-reactivity, we compiled five TCR databases containing high-resolution(*74–78*), sample-specific HLA information, totaling 1,603 samples (1,405 with complete class I and class II and 198 with only class I HLA data, Fig. 1). We performed TCR cluster-based enrichment that incorporates all TCRs to maximize the statistical power for discovering HLA-associated TCRs (hereafter referred to as HA TCRs). Specifically, the top 30,000 most highly expanded TCRs from each sample were selected and pooled, resulting in a combined dataset of 46,963,799 β-chain TCRs. These TCRs were clustered using the ultra-fast TCR clustering algorithm GIANA(*73*), which groups TCRs based on similarity of the Complementarity Determining Region 3 (CDR3) and TCR variable gene. TCRs within the same cluster are likely to share antigen-specificity and HLA restriction(*73, 79, 80*). We then performed TCR cluster-based enrichment analysis on appropriately sized clusters and identified 9,125,142 significant TCR-HLA pairs (FDR < 0.05, Fisher’s exact test), which constitutes by far the largest dataset on this topic. We developed computational models THNet to leverage this large-scale TCR-HLA datasets for both prediction and discovery. THNet can be applied to any high-throughput bulk βTCR sequencing sample for sample HLA typing prediction, offering broad HLA allele coverage with high accuracy. Furthermore, the HLA functional similarity network was utilized to perform population clustering and to assess the HLA mismatch tolerance for a given donor-recipient pair across different transplantation types.

**Fig. 1.**
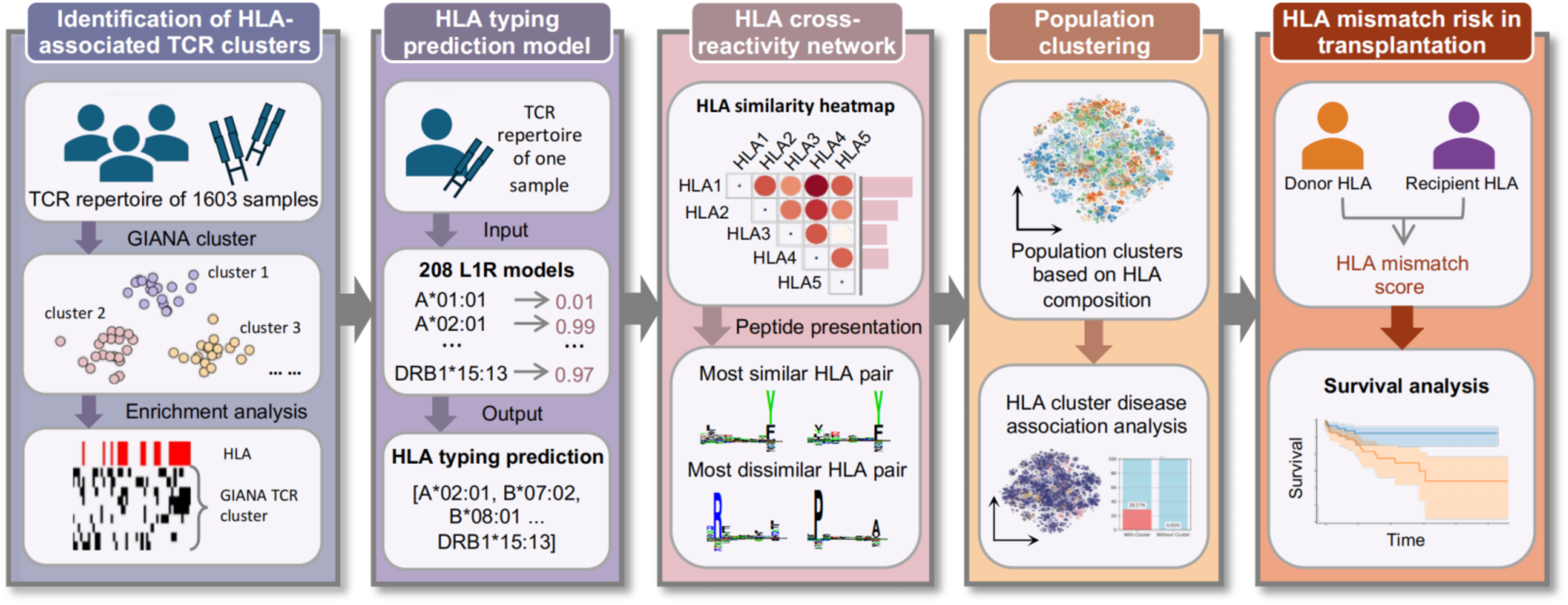
TCR-based HLA similarity mapping network. A schematic illustrating the THNet framework and its diverse applications. In this study, THNet was applied to two distinct bulk TCR-seq datasets to evaluate its HLA inference capability. Additionally, large-scale biomedical datasets and transplantation datasets were utilized to validate the HLA pair distance defined by THNet.

### Identification of HLA-Associated TCR clusters

The large number of HLA-associated TCR clusters allowed us to investigate TCR-HLA cross-reactivity. Notably, the number of HA TCRs varies greatly across different HLA alleles. More HA TCRs tended to be identified in alleles with a larger training sample size (Fig. 2A), yet the trend was insignificant (*p*-value =0.053) for class I alleles. This suggested that other factors, such as varying levels of TCR-HLA cross-reactivity, may influence the number of HA TCRs associated with each HLA allele. Despite this variability, we uncovered an average of 10,000 enriched TCRs for each allele.

**Fig. 2.**
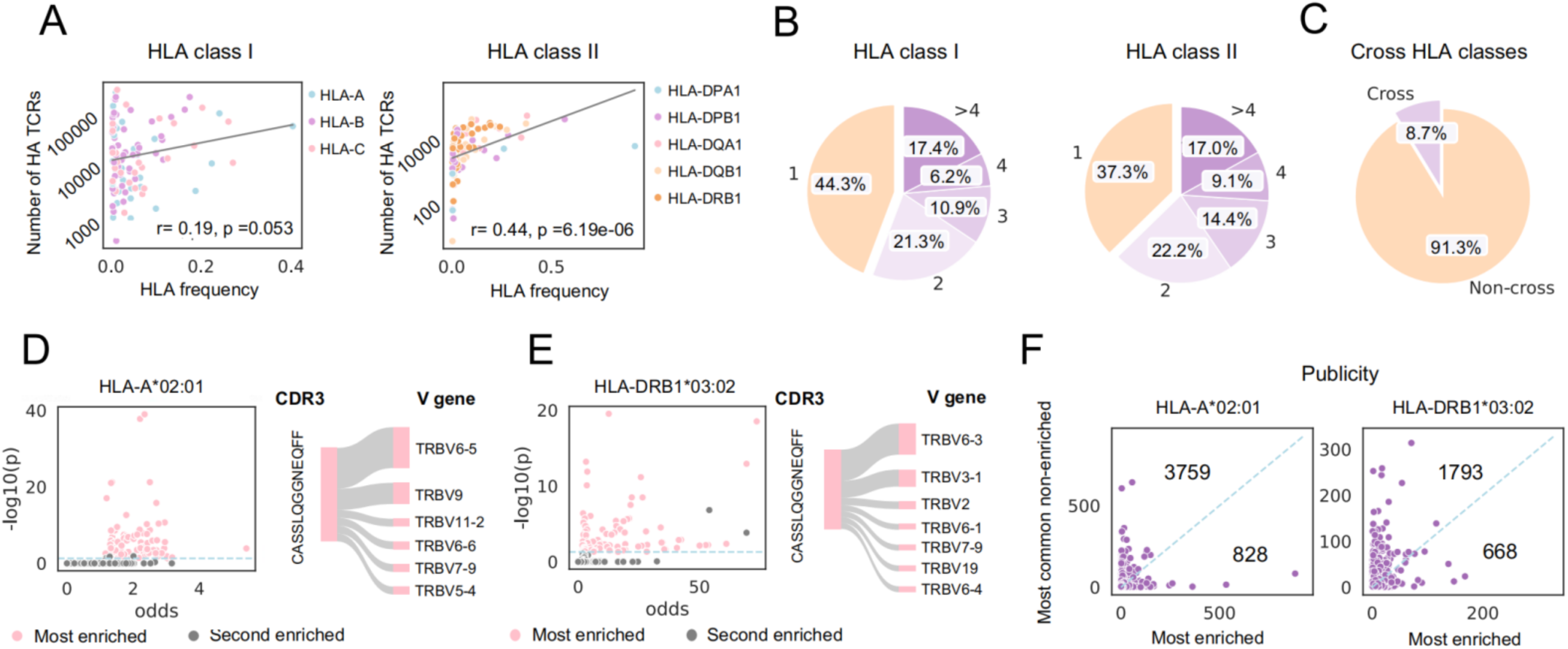
Identification of HLA-associated TCR clusters. (A) The number of HA TCRs for class I and class II HLA alleles, with each dot representing an HLA allele. (B) The ratio of HA TCRs enriched in n (where n = 1, 2, 3, 4, or >4) HLA alleles. (C) The number of HA TCRs cross-enriched across class I and class II HLA loci. (D–E) The odds ratio and negative log₁₀-transformed p-value of CDR3 regions of HA TCRs paired with different V genes. The pink dot indicates the most enriched CDR3–V gene combination, while the grey dot indicates the second most enriched combination. The Sankey diagram illustrates how a CDR3 displays distinct enrichment levels when paired with different V genes, with the width of each V gene path representing the level of enrichment. Panel (D) shows an example of HLA-A*02:01 (class I HLA), and panel (E) shows an example of HLA-DRB1*03:02 (class II HLA). The light blue dashed line represents the threshold where *p* = 0.05. (F) The publicity of the most enriched CDR3–V gene combination (x-axis) compared to the most prevalent combination among the remaining V genes (y-axis), shown here with examples of HLA-A*02:01 and HLA-DRB1*03:02 representing class I and class II HLA, respectively. Publicity is defined as the number of individuals in the training data who possess this TCR. The blue dashed line represents where y=x.

Most HA TCRs were associated with more than one allele (Fig. 2B). Specifically, only 44.3% of class I and 37.3% of class II HA TCRs were enriched in a single HLA allele, while 17.4% of class I and 17.0% of class II HA TCRs were enriched in more than four HLA alleles. This finding suggests a high level of TCR-HLA cross-reactivity, where a single TCR can be selected by multiple HLA alleles. However, TCRs enrichment across both class I and class II HLA alleles were much less common compared to cross-reactivity within the same HLA class (Fig. 2C), consistent with the distinct maturation paths of CD4⁺ and CD8⁺ T cells.

Next, we investigated whether HLA restriction is determined by the CDR3 region, the V gene, or the combination of both. We paired the CDR3 regions of HA TCRs with various V genes and compared the enrichment levels of different CDR3–V gene combinations. Nearly all CDR3 regions exhibited significant enrichment only when paired with a specific V gene (Fig. 2D-2E). To determine whether the most enriched CDR3–V gene pairs resulted from their population prevalence, we compared the publicity of the most enriched pairs with remaining V genes combination. The results showed that, in most cases, the most enriched CDR3–V gene pair was not the most prevalent combination for that CDR3 (Fig. 2F), confirming that the observed enrichment is driven by the interaction between TCR and HLA, rather than TCR publicity. Similarly, V genes exhibited HLA enrichment only when paired with specific CDR3 regions (Fig. S1A). These findings indicate that HLA restriction is determined by the specific combination of both the CDR3 region and the V gene, rather than by either component alone.

### Reproducible HLA typing inference from TCR Repertoire

To systematically characterize TCR-HLA interactions, we developed **THNet**, a framework leveraging 9 million significantly associated HLA-TCRs. Before exploring the broader implications of TCR-HLA cross-reactivity, we first assessed whether HLA alleles could be inferred from an individual’s TCR repertoire. This predictive capability serves as an initial validation that TCR repertoires contain distinct HLA-specific patterns. To do this, we built an L1 regularized logistic regression model for each HLA allele using their associated TCRs as the input features (see Methods).

One of the primary challenges in constructing HLA inference models is the vast diversity of HLA alleles. Aside from a few common alleles like HLA-A*02:01, most HLA alleles have low population frequencies(*81, 82*), resulting in limited positive samples in the training data. Our TCR cluster-based approach identified significantly more HA TCRs for each HLA allele than a previous method HLAGuessr(*83*) (Fig. S1B), enabling broader HLA coverage. In total, THNet can predict 109 class I HLA alleles and 99 class II HLA alleles. We calculated the AUC for 194 HLA models (with at least one positive sample in the testing data), revealing that 91% of class I and 86% of class II HLA models achieved an AUC greater than 0.8 (Fig. 3A). Additionally, the F-scores of all 208 HLA alleles were calculated using the entire combined dataset (Fig. S2A), with 95.4% of class I and 91.9% of class II HLA alleles exceeding an F-score of 0.8. As expected, the F-score of each HLA allele model positively correlated with the number of positive samples (*R* = 0.21, *p* = 0.0022; Fig. S2B). To further evaluate HLA coverage, we compared how many HLA alleles were covered per sample between THNet and HLAGuessr across datasets from three distinct regions (Hong Kong, Mexico, and Russia) with population-specific HLA compositions. THNet consistently covered significantly more HLA alleles across all tested datasets (Fig. 3B).

**Fig. 3.**
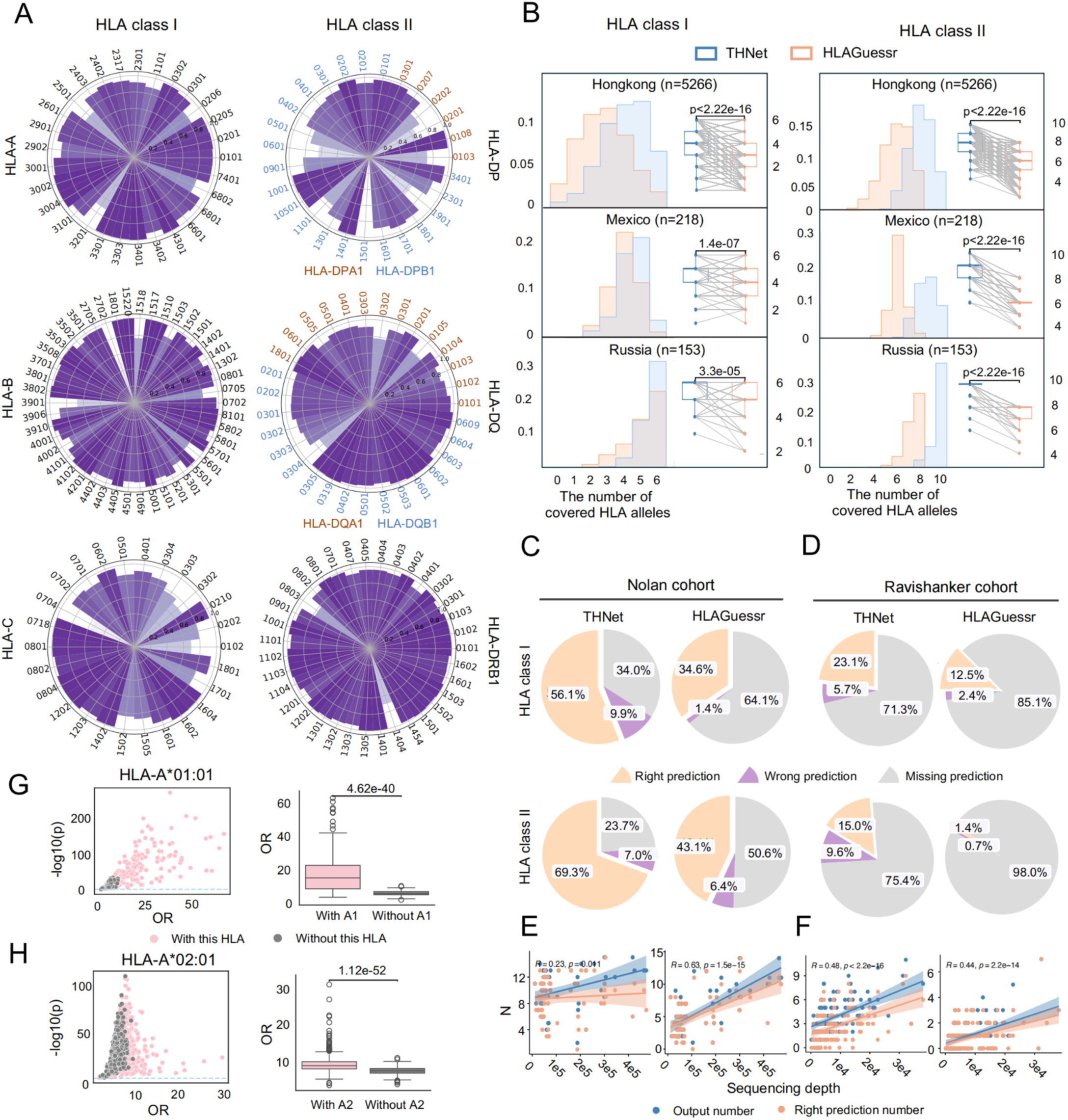
Reproducible HLA inference from TCR repertoire. (A) Polar bar chart displaying the AUC of each HLA allele model tested on the test dataset. Darker colors represent values closer to 1, while lighter colors indicate values closer to 0.5. The alpha and beta chains of HLA-DP and HLA-DQ are colored in orange and blue, respectively. (B) Comparison of HLA coverage between THNet and HLAGuessr across populations from different regions (Hong Kong, Mexico, and Russia). The left panel shows a bar plot of the density distribution of how many HLA alleles present in a sample are covered in the predictable HLA allele list. The right panel presents a box plot comparing the number of covered HLA alleles per sample between THNet and HLAGuessr, where each dot represents a sample, and the same sample is connected by a grey line across the two models. In both plots, THNet is shown in blue and HLAGuessr in orange. (C-D) Pie charts showing the ratio of right, wrong, and missing predictions for THNet and HLAGuessr on the Nolan and Ravishankar datasets, respectively. (E-F) Scatterplots illustrating the association between the number of all predicted outputs or correct predictions (y-axis) and sequencing depth (x-axis) for THNet and HLAGuessr on the Nolan and Ravishankar datasets. The total predictions and correct predictions are colored in blue and orange, respectively. Each dot represents a single sample, with consistent model and dataset annotations as in panels (C-D). The 95% confidence interval is indicated. (G–H) Plots showing the enrichment levels of HLA-restricted antigen-specific TCRs among HA TCRs. The left scatterplot displays the odds ratio (x-axis) and negative log₁₀-transformed p-value (y-axis) from Fisher’s exact test for each sample in the Nolan cohort. The light blue dashed line represents the threshold where p = 0.05. The right box plot compares the odds ratios of this enrichment between samples with and without the specific HLA allele. Panel (G) shows an example for HLA-A*01:01, while panel (H) shows an example for HLA-A*02:01.

We next compared the performance of THNet and HLAGuessr using two independent datasets. The first, the Nolan cohort, included 63 samples with an average of 132,101 unique TCR reads(*84*). The second, the Ravishankar cohort, consisted of 134 samples with an average of 8,607 unique TCR reads(*85*). Both datasets provided four-digit HLA information, although the Nolan cohort only included two-digit resolution for class II HLA alleles. THNet demonstrated comparable accuracy while producing significantly more right predictions (Fig. 3C–3D). Although the performance of both models was influenced by sequencing depth, THNet remained relatively robust when TCR sequencing exceeded 100,000 reads (Fig. 3E–3F).

Additionally, we discovered that HA TCRs were more likely to be HLA-restricted antigen-specific TCRs. Using THNet, we inferred the HLA types of another 1,414 samples in the Nolan cohort lacking HLA allele information, selecting only high-confidence predictions. Fisher’s exact test was then used to assess the enrichment of HLA-restricted antigen-specific TCRs within HA TCRs (see Methods). Results showed significant enrichment in nearly all tested samples—99.5% for HLA-A*01:01 and 100% for HLA-A*02:01 (Fig. 3G–3H). Furthermore, the odds ratio (OR) between these TCR groups was substantially higher in individuals carrying the corresponding restricted HLA allele compared to those who did not (Fig. 3G–3H). These findings offer new insights into the discovery of antigen-specific TCRs. Taken together, the high accuracy of our predictive model and the broad HLA coverage achieved through TCR cluster-based enrichment collectively underscore the validity of using this large-scale TCR-HLA association dataset to decode immune recognition patterns. These results highlight the structured nature of TCR-HLA interactions across independent individuals and demonstrate that TCR β chain-only data is sufficient for capturing meaningful immune relationships. This predictive framework provided the rationale to investigate HLA-restricted TCR repertoire and their disease associations.

### TCR-based HLA similarity network

Next, we sought to investigate whether different HLA alleles can select a shared fraction of the TCR repertoire, an indication of potential cross-reactivity. To explore, we constructed an HLA functional similarity network using HA TCRs. Each TCR was transformed into a numerical embedding using our previously developed method(*34*), and the Hausdorff metric was used to calculate the mismatch level between every pair of HLA alleles as their distance (see Methods). The resulting distances were normalized to a 0–1 scale (Fig. 4A–4B), where higher values indicate greater dissimilarity of the two TCR repertoires and, consequently, lower degree of cross-reactivity. Our analysis revealed that class I HLA alleles exhibit higher cross-reactivity with each other compared to class II HLA alleles. The average distance among class I HLA alleles was 0.72, while the average distances for class II HLA alpha and beta chains were 0.79 and 0.83, respectively.

**Fig. 4.**
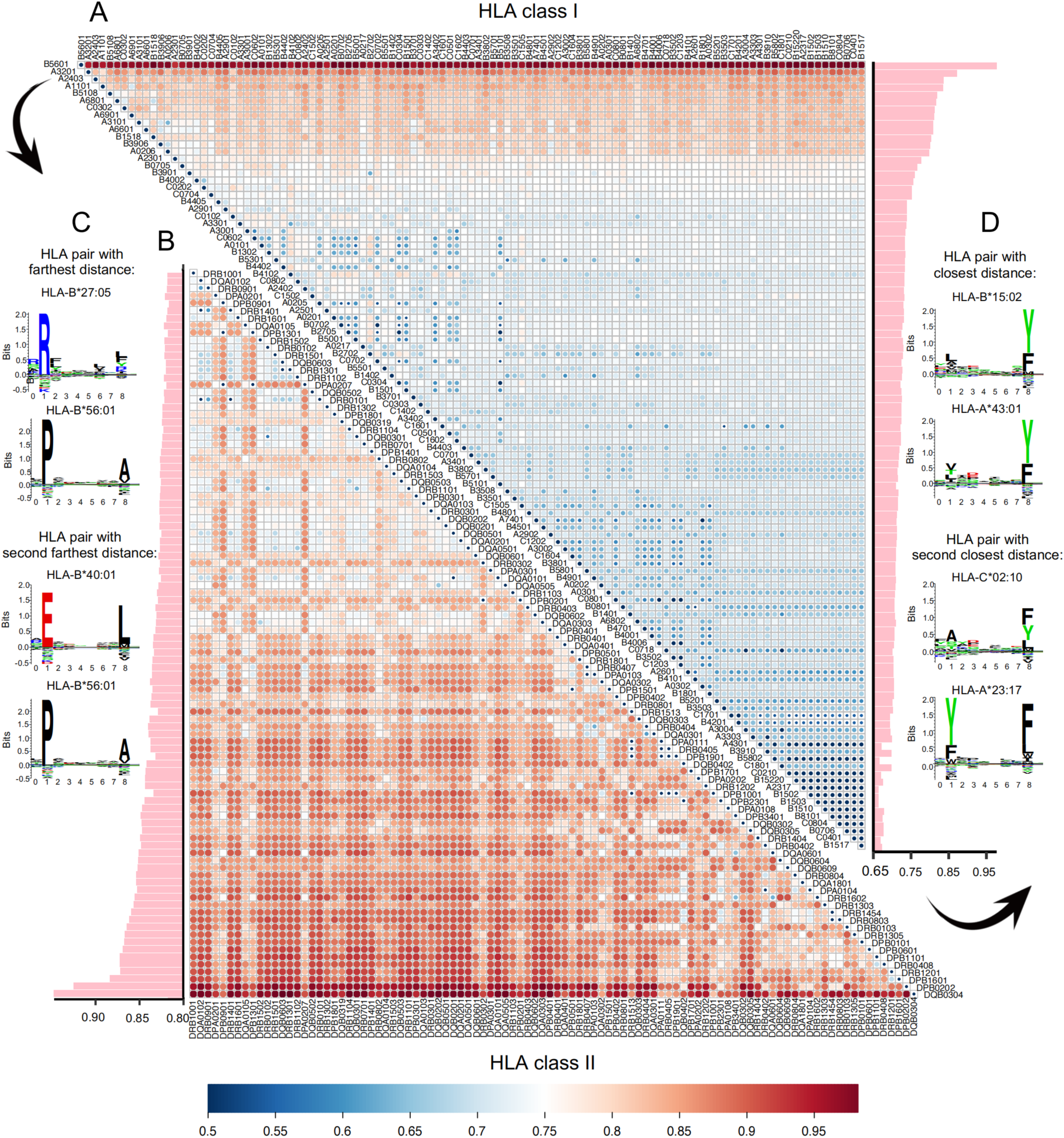
TCR-based HLA similarity network. (A–B) Heatmaps displaying the pairwise distance between class I and class II HLA alleles, respectively. The color gradient transitions from blue to red as the distance value increases. Pink bar plots adjacent to the heatmaps indicate the average distance between each HLA allele and all other alleles. To enhance contrast, values less than 0.5 were set to 0.5 in the heatmaps. The raw data are available in the supplementary materials. (C) Motifs of two class I HLA pairs (HLA-B*27:05 and HLA-B*56:01; HLA-B*40:01 and HLA-B*56:01) exhibiting the greatest distance. (D) Motifs of two class I HLA pairs (HLA-B*15:02 and HLA-A*43:01; HLA-C*02:10 and HLA-A*23:17) exhibiting the smallest distance.

We further investigated the relationship between HLA distances and their peptide-binding capacities. To begin, we explored the peptide-binding motifs of the two most dissimilar and two most similar HLA pairs (Fig. 4C–4D). Peptide-binding motifs were obtained from naturally presented ligands predicted using the NetMHCpan-4.1 online server(*86*). The most dissimilar class I HLA pair, HLA-B*27:05 and HLA-B*56:01, exhibited distinct peptide-binding features. Arginine was the dominant anchor AA at position 1 for HLA-B*27:05, whereas proline dominated the same position for HLA-B*56:01. These two AAs differ significantly in structure, chemical properties, and biological functions, leading to diverse peptides presented by these HLA alleles and, consequently, distinct TCR repertoires. Similarly, the second most dissimilar pair, HLA-B*40:01 and HLA-B*56:01, displayed exclusive peptide-binding motifs at anchor position 1 (Fig. 4C). In contrast, the two most similar HLA pairs, HLA-B*15:02-HLA-A*43:01 and HLA-C*02:10-HLA-A*23:17, shared highly similar peptide-binding motifs without contradictory dominant anchor AAs (Fig. 4D).

To validate these individual observations on a larger scale, we analyzed a dataset of 139,020 experimentally validated peptide-HLA pairs from the Siranush cohort(*87*) that met the prediction criteria of NetMHCpan-4.1(*86*) (see Methods). For each peptide-HLA pair, we replaced the HLA allele with its closest and farthest counterparts and predicted peptide-binding affinity using NetMHCpan-4.1. Peptides presented by a given HLA allele were found to have significantly higher binding affinities with its close counterparts than with more distant ones (Fig. S3A). There is also higher proportion of peptides binder among close HLA counterparts compared to those with greater distances (Fig. S3B). These results demonstrate that the HLA similarity network, defined based on their TCR outputs, reflects the similarity of peptide presentation across HLA alleles.

### HLA-based population clustering and disease association

We utilized our HLA similarity network to cluster populations into distinct groups based on their class I and class II HLA allotypes. HLA alleles are putative risk predictors for autoimmune disorders, such as HLA-DRB1*03-DQB1*0201 in Type-I Diabetes and HLA-B27 in Ankylosing spondylitis*^(88–91)^*. By grouping functionally similar alleles into clusters, we may increase the statistical power to identify new disease relevance. In this analysis, we took advantage of approximately half of a million samples from the UK Biobank database(*92*). We first selected samples with complete HLA information and then grouped identical HLA genotypes together, resulting in 65,323 and 36,597 unique class I and class II allotypes, respectively. The sum of pairwise distances between HLA allotypes were calculated as the correlation of one allotype relative to all others in the dataset. For class I HLA clustering, this process generated a 65,323 × 65,323 matrix (and a 36,597 × 36,597 matrix for class II HLA clustering), which was visualized in t-SNE plots (Fig. 5A, Fig. S4A). The samples from different ethnic backgrounds are well-separated, indicating distinct HLA allele distributions in different populations. We selected three major populations (White, Asian, and Black) for further analysis to avoid race-driven biases in the downstream analysis. Individuals with functionally similar HLA genotypes were grouped into 340, 140, and 140 clusters for White, Asian, and Black populations, respectively. Disease enrichment for each cluster were evaluated using Fisher’s exact test, and p-values were adjusted for multiple testing using the Benjamini and Hochberg false discovery rate correction.

**Fig. 5.**
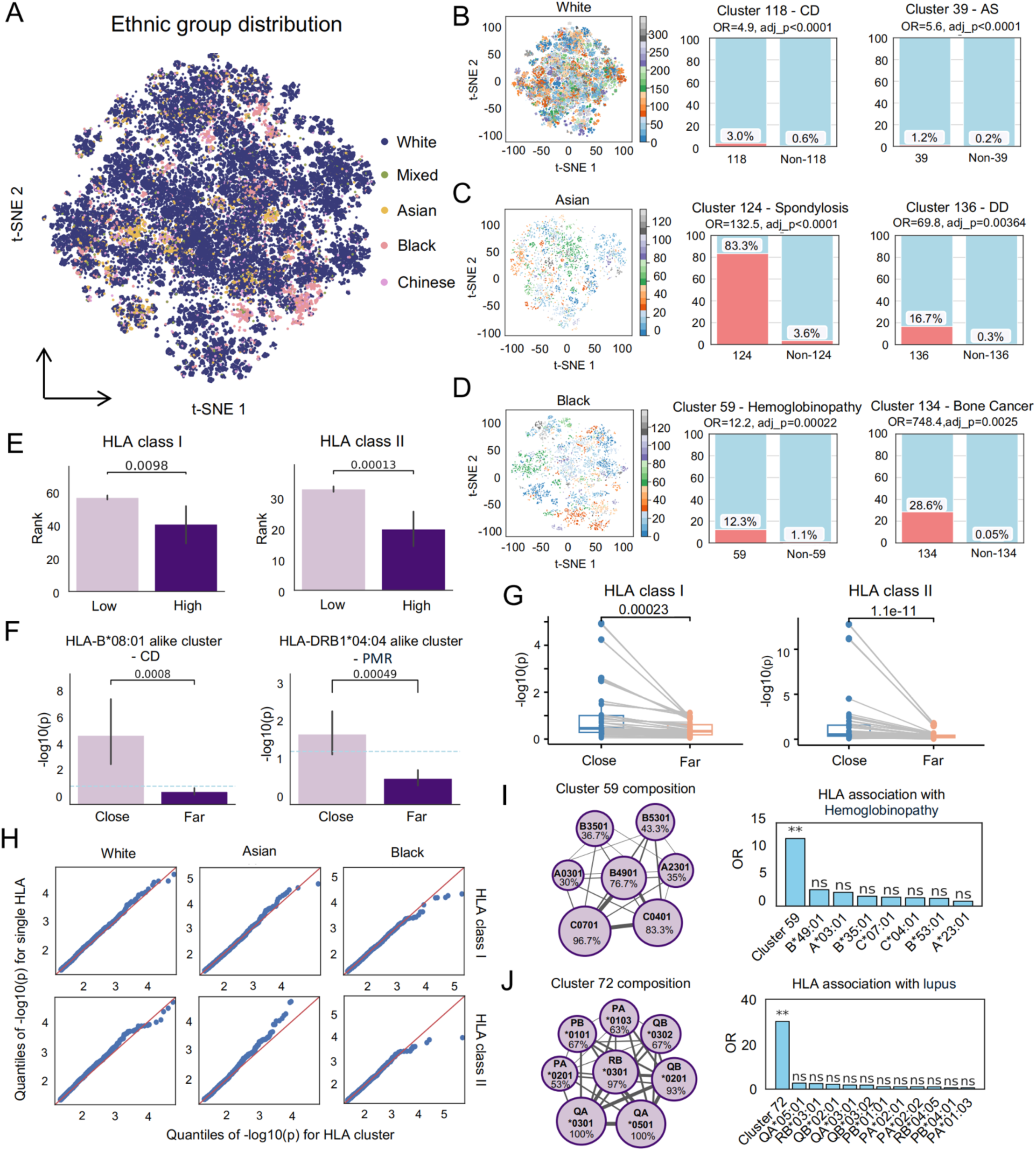
HLA-based population clustering. (A) t-SNE plot depicting population clustering in the UK Biobank dataset based on class I HLA composition. Samples are colored according to race annotations in the UK Biobank database. Each dot represents one or more samples sharing the same HLA class I genotype. If a genotype is shared by samples from different racial backgrounds, the color is assigned based on the most frequent race within that genotype. (B–D) t-SNE plots displaying population clustering for White, Asian, and Black British populations, accompanied by stacked bar plots showing Fisher’s exact test results for the two clusters with the most significant disease associations within each population. The x-axis of the bar plots indicates samples with/without the HLA cluster, while the y-axis represents the percentage of samples with specific diseases (CD: celiac disease; AS: ankylosing spondylitis; DD: developmental delays). (E) Bar plot comparing the distance rank of HLA alleles with high or low disease associations similarity relative to specific HLA alleles. (F) Examples of disease associations for HLA clusters close to or far from disease-associated HLA alleles. The y-axis represents the negative log10-transformed p-value of the Fisher’s exact test for disease associations with each HLA cluster. The left example shows CD associations for clusters close to or far from HLA-B08:01, a known CD risk allele. The right example illustrates polymyalgia rheumatica (PMR) associations for clusters close to or far from HLA-DRB1*04:04, a known PMR risk allele. The light blue dashed line represents p = 0.05. (G) Boxplots comparing the enrichment levels of HLA clusters close to or far from disease-risk HLA alleles. Each dot represents a disease-risk HLA allele pair, with the same pair connected by grey lines between the close and far HLA cluster groups. The y-axis represents the average enrichment level of each HLA cluster group, with the groups defined based on whether their distance to the risk allele is above or below the median value. (H) Quantile–quantile (QQ) plots comparing the quantiles of negative log10-transformed p-values for HLA-disease associations. The y-axis represents results from single-HLA enrichment, and the x-axis represents results from the HLA cluster-based approach. The maximum value of x-axis and y-axis of White population were restricted to 5 to better display the majority of the sample points. The red line represents where y=x. (I–J) Composition and disease associations of class I and class II HLA clusters 59 and 72 in black population, respectively. Disease associations are shown for both the clusters and their composing HLA alleles. In the network plots, circle size and the percentage in the circle represent the frequency of an HLA allele within the cluster, while line width indicates the frequency of co-occurrence between two HLA alleles. For simplicity, only HLA alleles that appear in over 50% of the samples in a class III HLA cluster are displayed. PA: DPA1; PB: DPB1; QA: DQA1; QB: DQB1; RB: DRB1. Significance levels: ‘**’ (adj p < 0.001), ‘*’ (adj p < 0.01), and ‘ns’ (not significant).

Significant disease associations of specific HLA clusters were identified in all tested populations, for both class I and class II HLAs (Fig. 5B–5D, Fig. S4B–S4D). For example, samples in cluster 118 exhibited a higher risk of celiac disease (CD) compared to those outside the cluster (Fig. 5B). All significant class I and class II HLA cluster disease associations for populations with different ethnic backgrounds are provided in the supplementary tables. The dataset included 416,967 White British samples, but only 5,615 Asian and 3,751 Black samples, leading to some notably high odds ratios (OR) in smaller populations, such as the association of cluster 134 with bone cancer in Black British samples (OR= 748.4, Fig. 5D). Due to the larger sample size, OR estimates for the White population are likely more robust and reflective of true disease associations. Consequently, most downstream analyses were focused on the White population.

We examined whether the HLA clusters were confounded by HLA haplotypes, where certain HLA pairs in linkage disequilibrium (LD) tend to be inherited together more frequently than expected. Our results indicate that nearly all tested individuals carry at least one HLA haplotype (98.8% for HLA class I and 100% for HLA class II loci, Fig. S5A). However, fewer clusters are predominantly composed of HLA alleles in LD (26.0% for class I and 46.6% for class II loci, Fig. S5A). Among the disease-associated HLA clusters, 29.1% of class I HLA clusters and 65.5% of class II HLA clusters are dominated by HLA haplotypes in LD (Fig. S5B). Notably, even within these LD-dominated disease-associated clusters, disease enrichment exhibited significantly higher odds ratios (ORs) compared to using HLA haplotypes alone (Fig. S5C). These results suggested that our TCR-based distance measure captures functional HLA similarities despite the presence of haplotypes.

We next defined two alleles to have high disease-association similarity if they share significant associations with more than five diseases. The distances between these HLA pairs were significantly shorter compared to pairs with low disease-association overlap (Fig. 5E). Additionally, individuals in the clusters with HLA genotypes closer to disease-risk alleles exhibited higher risks for the associated diseases, where a “disease-risk HLA allele” was defined as the allele most significantly associated with a disease in the dataset. The distance between an HLA cluster and a risk HLA allele was calculated as the mean weighted sum of pairwise distances (see Methods). Clusters closer to risk HLA alleles, such as HLA-B*08:01 for CD or HLA-DRB1*04:04 for polymyalgia rheumatica (PMR), showed higher disease enrichment when compared to single HLA alleles (Fig. 5F). This observation was generally true to both class I and class II alleles (Fig. 5G).

Further, we found that advantage of HLA cluster over single allele in disease association was more obvious among the Black population. While the p values of single-HLA enrichment and HLA cluster-based approaches showed similar distributions in White and Asian populations, cluster-based approach identified more significant HLA-disease associations in Black people (Fig. 5H). This may be attributed to the shorter LD blocks in individuals of African ancestry(*93*), making cluster-based approaches more effective in grouping functionally relevant allelic combinations and capturing meaningful disease associations. For example, the HLA compositions of class I cluster 59 and class II cluster 72 exhibited the strongest disease associations in the Black population (Fig. 5I-J). They were enriched for hemoglobinopathy and lupus respectively. Notably, none of the composing HLA alleles within these clusters showed significant enrichment (Fig. 5I-J). In summary, the use of HLA clusters defined by the HLA distance matrix enables the identification of disease associations at the whole HLA genotype level, uncovering novel HLA disease susceptibilities that that cannot be detected through single-allele analyses.

### HLA mismatch score as an indicator of transplantation outcome

We next extended the HLA similarity analysis to quantitatively assess HLA mismatches during organ transplantation. Fewer HLA mismatches between donor and recipient is generally associated with better transplantation outcomes and longer survival(*94–96*). Further, the clinical impact of mismatches is depending on the similarity between HLA alleles, where mismatches involving more similar HLA alleles are typically more tolerable than those between highly distinctive alleles(*96–98*). Eplet score is the most commonly used tool for HLA similarity evaluation, which quantifies differences in surface amino acids of HLA molecules between donors and recipients(*98–100*). It assumed that mismatched HLA alleles with greater structural differences are more likely to elicit donor-specific B cell responses that may lead to graft failure(*100–102*). In contrast, HLA similarity defined in our study is based on its associated TCR repertoire, which might complement the B-cell-focused eplet method, and provide an orthogonal perspective on the role of HLA mismatch in transplantation outcomes.

To explore, we analyzed the Scientific Registry of Transplant Recipients (SRTR) database(*103*), which has 923,533 recorded entries on donors and recipients of organ transplantation in the United States since 1987. 84,816 entries with HLA genotypes of both donors and recipients, as well as transplantation outcomes, were included in the analysis. Since high-resolution HLA alleles were unavailable for most individuals in this dataset, we imputed 4-digit HLA alleles from two-digit records within specific populations (see Methods). Samples containing unpredictable HLA alleles were excluded from subsequent analysis. Since patient outcomes were significantly associated with the race of both donors and recipients (Fig. S6A–S6B), we focused our analysis on the White population due to its largest sample size.

We first studied kidney transplant cases given their highest abundance among transplantation types, with 3,234 passed-filter entries. The THNet mismatch score (MS) was calculated as the sum of the mean pairwise distances against recipient for each donor-specific HLA allele, separately for class I and class II (Fig. 6A, see Methods). THNet MS was independent of the eplet score when conditioned on the number of HLA mismatches (Fig. S7A–S7B). Class I MS demonstrated superior predictive power for kidney graft failure compared to the eplet, while the case was the opposite for class II (Fig. 6B). We thus combined the THNet class I MS and the eplet class II score by selecting the greater value of the two. The new predictor provided optimal stratification of patients based on the timing of graft failure (Fig. 6C). Additionally, the combined score displayed a higher hazard ratio (HR = 1.33, *p* < 0.001) compared to eplet alone (HR = 1.04, *p* = 0.005) when adjusted for potential confounders such as HLA mismatch number (MN), age, BMI, and gender (Fig. 6D).

**Fig. 6.**
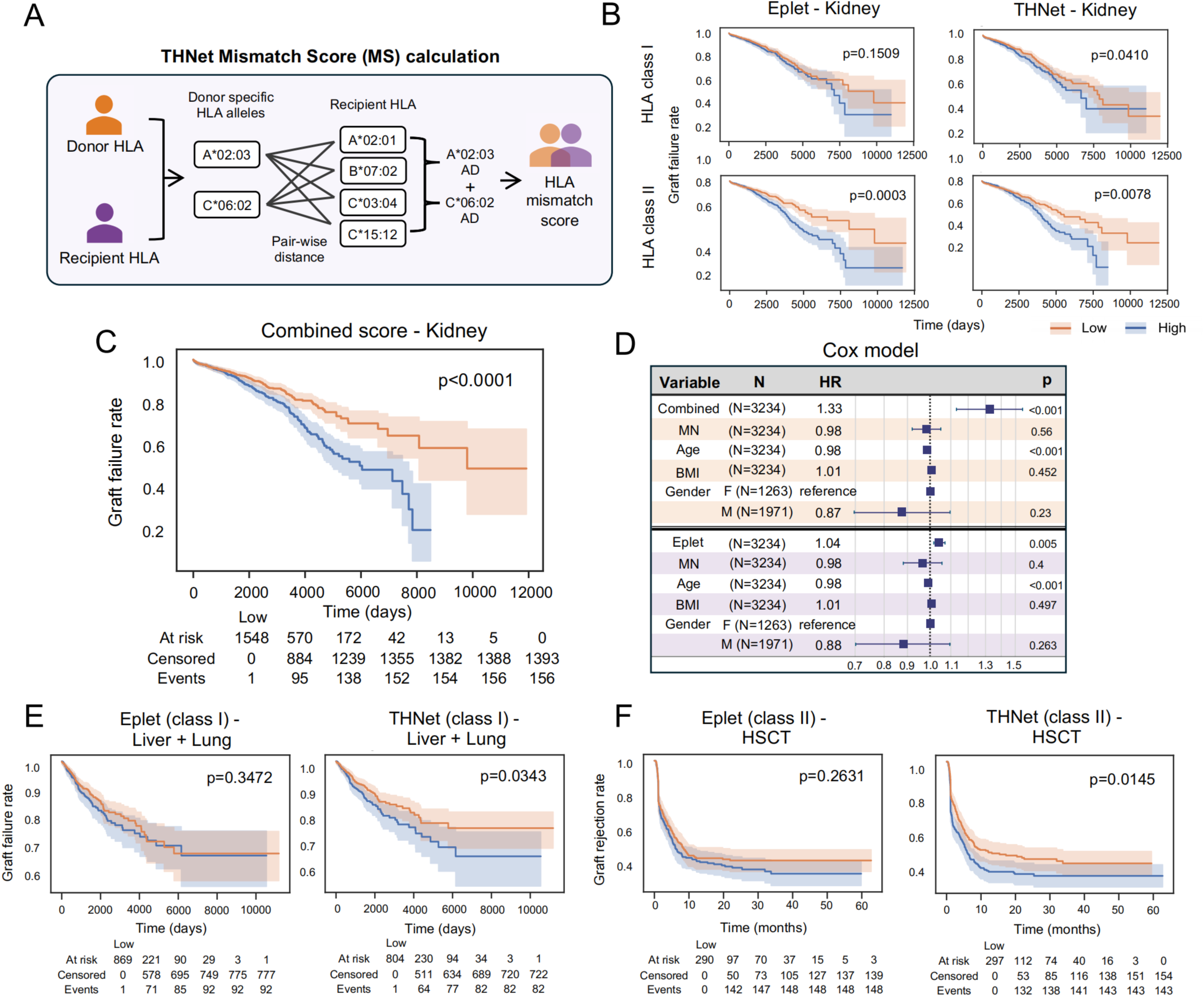
HLA mismatch score as an indicator of transplantation outcome. (A) Illustration of how the THNet mismatch score (MS) is calculated. (B) Kaplan-Meier curves showing kidney transplant failure rates for patients grouped by low/high eplet scores and THNet MS, with results for class I and class II HLA scores presented in two rows. High-value groups in the survival analysis are defined as individuals with scores above the median value. Statistical significance was assessed using the log-rank test. (C) Kaplan-Meier curve depicting graft rejection rates of kidney transplant patients using the combined score as an indicator. (D) Cox proportional hazards model showing hazard ratios and *p*-values for two models testing the combined score and eplet score as risk factors, adjusted for potential confounders, including HLA allele mismatch number (MN), age, body mass index (BMI), and gender. F: female; M: male. (E) Kaplan-Meier curves showing liver and lung transplant failure rates for patients grouped by low/high class I eplet scores and class I THNet MS, with the number of events indicated throughout the follow-up period. (F) Kaplan-Meier curve showing hematopoietic stem cell transplantation (HSCT) rejection rates for patients grouped by low/high class II eplet scores and class II THNet MS, with the number of events shown throughout the follow-up period.

We then assessed whether the THNet MS could be applied to other types of organ transplantation. Lung and liver transplant data were combined into a single cohort of 1,612 pairs from the SRTR dataset to increase power. Consistent with kidney transplant findings, the THNet class I MS displayed significantly better predictive power for graft failure than eplet score (Fig. 6E), while neither approach’s class II mismatch score was significant (Fig. S8C). To this end, THNet class I MS was the only indicator consistently associated with patients’ survival across all organ transplantation types investigated (Fig. S8A–S8B).

Lastly, we tested the applicability of the THNet in hematopoietic stem cell transplantation (HSCT) using a public cohort in the CIBMTR database(*104*), which contained high-resolution HLA typing for both donors and recipients. A total 594 transplantation entries passed our selection criteria (Methods). Interestingly, THNet class II MS (*p* = 0.0145) was a better predictor of graft rejection compared to the class II eplet score (*p* = 0.2631, Fig. 6F), while the class I score of neither method demonstrated an association with graft rejection rates (Fig. S8D). Overall, these findings highlight the utility of the THNet MS as a complementary tool to eplet, providing improved stratification of patients and enabling a more comprehensive assessment of donor-recipient immune compatibility across different transplantation settings.

## DISCUSSION

In this study, we conducted a large-scale investigation of HLA-wide cross-reactivity of T cell receptors by leveraging an extensive dataset of over 9 million TCR-HLA pairs. The reproducibility of the TCR cross-reactivity was successfully validated in unseen samples. Based on this result, we quantitatively estimated the shared TCR repertoires of different HLA alleles and defined the HLA similarity network. This network enabled the discovery of novel HLA-disease susceptibilities in the UK Biobank cohort. Additionally, mismatch score defined by HLA distances were shown to be a novel predictor of graft rejection in both organ transplantation and HSCT.

In our analysis, we observed distinct performance between class I and class II THNet MS across organ transplantation and HSCT, likely due to the differing mechanisms of graft rejection. In organ transplantation, T-cell-mediated rejection is primarily driven by a host-versus-graft response, where mature CD8⁺ T cells serve as the key effectors attacking the transplanted organ(*105, 106*). In contrast, HSCT rejection is predominantly a graft-versus-host response, in which donor CD4⁺ T cells must first be educated by the host’s thymic cells before initiating a broader immune response against host tissues(*107, 108*). As a result, class I mismatches may have a more significant impact in organ transplantation, whereas class II mismatches could be more detrimental in HSCT, particularly in the context of T-cell-mediated rejection.

Defining the HLA restriction of a given TCR is challenging because TCR recognition involves both HLA and the antigen peptide. Conventionally, the assessment of HLA restriction for a given TCR would require binding experiments against all available HLA alleles and the entire peptide pool, which is nearly impossible. Here, we defined TCR-HLA associations based on the significant co-occurrence patterns of TCR clusters among individuals with particular HLA alleles, akin to taking a glimpse at the results of “binding test” performed by the immune system *in vivo*. The resulting TCR-HLA associations do not represent the absolute HLA restriction of each TCR but rather reflect the outcomes of immune selection shaped by real-life antigen encounters. This approach may provide even more relevant insights in disease contexts, as it emphasizes functionally relevant immune interactions over theoretical HLA restrictions.

Among the many attempts to group similar HLA alleles together(*59, 60, 109–111*), our method is unique in defining similarity between HLA pairs not based on the allele sequences or structures, but rather on their functional outcome – the associated TCR repertoire. Indeed, sequence similarity is often associated with functional similarity. However, the TCR-pHLA interaction is highly sensitive, as a single AA replacement can disrupt the entire binding process, making it more susceptible to subtle changes(*112–114*). Given the limited understanding of how specific AA replacements at different positions within the HLA molecule impact this interaction, directly linking sequence similarity to functional similarity in this context may be problematic. Therefore, inference based on the functional outcomes might be more biologically relevant.

There are several limitations of this work. First, the number of predictable HLA alleles is still limited compared to the vast diversity of HLA alleles identified so far. Expanding datasets with high-resolution HLA genotypes is crucial for increasing the size of the HLA allele pool. Second, the identified TCR-HLA pairs may be influenced by the presence of HLA haplotypes. While the observed association is actually driven by one HLA allele, the other alleles in LD may also be labeled as associated with the same TCR. Third, it is likely that not all TCRs within a single cluster share the same HLA restriction. Misclustering TCRs specific to different HLA alleles will compromise the statistical power in our downstream analysis, yet will unlikely introduce false positive associations. Lastly, this study was conducted using bulk TCRβ chain sequencing datasets without knowledge of their paired ɑ chains. The lack of pairing limits the ability to fully characterize TCR-HLA interactions and may lead to an overestimation of TCR-HLA cross-reactivity. While single-cell TCR (scTCR) sequencing enables chain pairings, current scTCR-seq samples are not sufficient for the size and scale required by our analysis. Future advances in scTCR sequencing technology with enhanced scalability and cost efficiency will be essential to overcoming this barrier and further refining our understanding of TCR-HLA interactions.

In summary, we presented THNet, a TCR-based HLA similarity mapping network built upon HLA-associated TCR clusters, which reveals previously unrecognized cross-reactivity patterns across HLA alleles. By leveraging TCR-HLA functional relationships, this approach enhanced our understanding of immune cross-reactivity, shedding light on disease susceptibility and graft rejection. Our findings highlighted the potential of THNet to advance precision medicine by elucidating the role of HLA alleles in shaping clinical outcomes.

## MATERIALS AND METHODS

### TCR clustering

In the first step, we complied five publicly available dataset with high-resolution HLA information, totaling 1,603 samples. Of these, 1,405 samples contained complete class I and class II HLA data, while 198 samples had only class I HLA data(*74–78*). Quality control (QC) was performed on the TCR sequences based on two criteria: (1) the TCR length must fall between 9 and 23 amino acids, and (2) the TCR CDR3 regions must start with Cysteine (C) and end with Phenylalanine (F). Following QC, we selected the top 30,000 most highly expanded T cell clones, as determined by clonal frequency, to reduce the computational burden of TCR clustering. The resulting TCR sequences were aggregated into a TCR pool comprising 46,963,799 unique sequences. To further reduce computational complexity, the TCR pool was randomly divided into 12 subgroups, each containing fewer than 4 million TCR reads. GIANA TCR clustering was then performed on each subgroup(*73*). To enhance the number of clustered TCRs, the TCR pool was re-shuffled three times, and the clustering process was repeated for each iteration. All resulting TCR clusters were subsequently integrated. To ensure the validity of downstream analyses, we applied additional filtering steps. Only clusters with fewer than 2,000 TCRs were retained, under the rationale that larger clusters are less likely to adhere to the assumption that TCRs within the same cluster share the same HLA restriction. Clusters that appeared in fewer than five samples were also excluded due to insufficient statistical power for HLA enrichment analysis. After filtering, 456,315 TCR clusters were retained, with an average of 10 TCRs per cluster.

### HLA enrichment analysis

We performed HLA enrichment analysis at the TCR-cluster level. A cluster was considered present in a sample if at least one TCR within the cluster was detected in that sample. If multiple TCRs from the same cluster appeared in a single sample, each occurrence was counted separately. The total occurrences for each cluster in relation to each HLA allele were then summed to represent the overall association value of the cluster with that allele. By aggregating all clusters, we constructed a matrix where the rows represented clusters, the columns represented HLA alleles, and each cell contained the occurrence value for a specific cluster-HLA allele pair. Next, we applied Fisher’s exact test to assess the enrichment of each cluster-HLA allele pair, using “greater” as the alternative hypothesis. The resulting p-values for all cluster-HLA allele pairs were corrected for multiple testing using the Benjamini-Hochberg false discovery rate (FDR) correction, performed separately for each HLA allele. Cluster-HLA allele pairs with an adjusted p-value less than 0.05 were considered significantly associated.

### The constriction of HLA typing inference models

The significantly associated TCRs for each HLA allele were utilized to build a separate logistic regression model with L1 regularization for each HLA allele. In these models, the presence of each TCR was assigned a weight, indicating how much the occurrence of that TCR contributes to the likelihood of the corresponding HLA allele being present. The rationale behind these models is to predict whether a sample contains a specific HLA allele based on the presence of its corresponding TCRs in the TCR repertoire. The probability formula for each HLA allele is as follows:

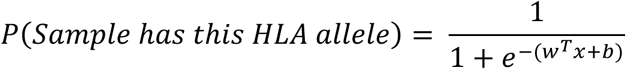

where *w* is the vector of learned coefficients for each TCR feature, *x* is the input feature vector (*x* =1 if the feature TCR is present, *x* =0 otherwise), and *b* is the bias term learned during training. The combined dataset was split into 80% for training and 20% for testing. Model was trained on training dataset using maximizing the log-likelihood, while the testing set was used to determine the optimal regularization strength. The thresholds for the final decision were determined based on the maximum F-score obtained from training on the entire combined dataset.

### The enrichment of HLA-restricted antigen-specific TCRs among HA TCRs

To test the association between HLA-associated (HA) TCRs and HLA-restricted antigen-specific TCRs, we analyzed SARS-CoV-2-specific TCRs from the Nolan cohort(*84*). The sample source of the TCRs used in the multiplexed identification experiment was available in the original study(*84*). All analyses in this section were conducted in an HLA allele-specific manner, with HLA-A*01:01 used here as an illustrative example. HLA-A*01:01-restricted antigen-specific TCRs were defined based on two criteria: (1) the TCRs originated from samples carrying the HLA-A*01:01 allele, and (2) the TCRs recognized peptides presented by HLA-A*01:01 molecules as predicted by predicted by NetMHCpan-4.1. We then applied Fisher’s exact test to assess whether HLA-A*01:01-restricted antigen-specific TCRs were significantly enriched among HLA-A*01:01-associated TCRs compared to non-HLA-A*01:01-associated TCRs in each sample. Furthermore, we used THNet to infer the presence of the HLA-A*01:01 allele in samples, selecting only those with high-confidence predictions (probability < 0.01 as HLA-A*01:01-negative and probability > 0.99 as HLA-A*01:01-positive). Finally, Welch’s t-test was used to compare the odds ratio of HLA-A*01:01-restricted antigen-specific TCR enrichment between HLA-A*01:01-positive and HLA-A*01:01-negative samples.

### The HLA similarity network

The HA TCRs were first converted into a numeric embedding using a method similar to the RFU approach(*34*), incorporating additional CDR1, CDR2, and CDR2.5 regions. The combined CDR1/2/2.5 region along with the CDR3 regions was each embedded into a 500-dimensional vector, which were then concatenated to form the final embedding for a TCR sequence. Each HLA allele was then represented by an n*1000 matrix, where n is the number of HA TCRs associated with that HLA allele and 1000 is the vector length of each TCR read. To quantify the distance between two HLA alleles, we used the Hausdorff distance(*115*), which is defined as:

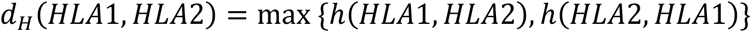

where *h*(*HLA*1, *HLA*2) computes the closest HLA1 TCR to any HLA2 TCR in the embedding space, and *h*(*HLA*2, *HLA*1) computes the closest HLA2 TCR to any HLA1 TCR in the embedding space. The Hausdorff distance between two HLA alleles is given by the maximum of these two values. The distances between all tested HLA alleles were computed separately for HLA class I and HLA class II. The resulting distance values were then normalized to the range of 0 to 1 using min-max normalization.

### Peptide binding pattern between HLA pairs with different distance

To compare the peptide binding patterns of HLA pairs with different distances, we utilized a dataset containing experimentally validated peptide binders for various HLA molecules. For each known HLA-peptide pair, we substituted the original HLA allele with a closely or distantly related counterpart and assessed its binding affinity to the original peptide using NetMHCpan-4.1(*86*). The rank scores (Rnk_EL) were used to quantify the binding ability of the tested HLA-peptide interactions, with a binding threshold set at 2% to classify peptides as HLA binders. Due to the limited predictive accuracy of NetMHCpan-4.1 for HLA class II alleles, this analysis was conducted exclusively for HLA class I alleles.

### Population clustering based on HLA composition

To cluster the population into smaller subgroups based on their HLA composition, we first computed a sample distance matrix, which is an n-by-n matrix, where n is the number of samples. Each value in the matrix represents the distance between a given sample and another sample. The sample-wise distance between two samples, *S_a_* and *S_b_*, is calculated as:

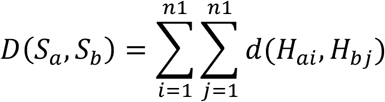

where *D*(*S*_*a*_, *S*_*b*_) denotes the overall distance between sample *S*_*a*_ and sample *S*_*b*_; *n*1 represents the number of HLA alleles per sample (*n*1=6 for class I HLA alleles and *n*1=10 for class II HLA alleles); *d*(*H*_*ai*_, *H*_*b*j_) is the distance between HLA allele *H*_*ai*_of sample *S*_*a*_and HLA allele *H*_*b*j_of sample *S*_*b*_, as defined by THNet. Thus, the sample-wise distance is computed as the sum of all pairwise distances between HLA alleles of the two samples. Once the sample distance matrix was obtained, we applied agglomerative hierarchical clustering to group similar samples. Each sample initially formed its own cluster, and clusters were iteratively merged based on their closest neighbors until the final clustering structure was achieved.

### Identification of HLA linkage disequilibrium (LD)

HLA haplotypes in LD were identified based on two criteria: (1) a ‘D’ metric greater than 0.5 and (2) an adjusted p value less than 0.05. The D value quantifies LD by measuring the non-random co-occurrence of two HLA alleles(116, 117). It is calculated as:

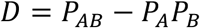

where P_AB_ is the observed frequency of the two HLA alleles A and B occurring together; and P_A_ and P_B_ are their respective marginal frequencies. To standardize *D*, we compute D′ as:

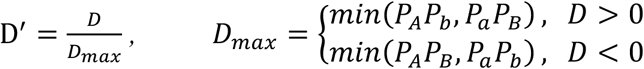

where *P*_*a*_=1-*P*_*A*_ and *P*_*b*_=1−*P*_*B*_ are the complementary allele frequencies. The D-value ranges from −1 to 1, with positive values indicating an increased co-occurrence of alleles A and B, and negative values indicating lower-than-expected co-occurrence. To ensure the robustness of identified HLA pairs in LD, we applied Fisher’s exact test, followed by false discovery rate (FDR) correction. Only allele pairs with a high D-value and a significant adjusted p-value were classified as HLA pairs in LD, while allele pairs with co-occurrence in fewer than five samples were excluded from the analysis. We defined an HLA cluster as LD-dominated if the two most frequent HLA alleles within the cluster is an HLA haplotype in LD.

### Distance calculation between HLA clusters and disease-risk HLA alleles

To quantify the distance between each HLA cluster and a disease-associated HLA allele, we first analyzed the disease association of all HLA alleles using Fisher’s exact test. After applying FDR correction, for each tested disease, the most significantly enriched HLA allele with an odds ratio > 2 was defined as a risk HLA allele. Next, we computed the distance between each HLA cluster and the identified risk HLA allele using the following formula:

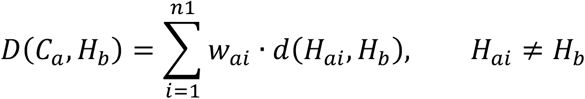

where *D*(*C*_*a*_, *H*_*b*_) denotes the overall distance between HLA cluster *C*_*a*_ and risk HLA allele *H*_*b*_, *n*1 represents the number of HLA alleles per cluster (only HLA alleles that appear in at least 25% of class I clusters and 50% for class II clusters were included), *d*(*H*_*ai*_, *H*_*b*_) is the HLA allele distance between HLA allele *H*_*ai*_in cluster *C*_*a*_ a and the risk HLA allele *H*_*b*_, as defined in THNet, *w*_*ai*_represents the frequency of allele *H*_*ai*_within cluster *C*_*a*_, serving as a weight in the calculation. If the risk HLA allele *H*_*b*_ is already present in HLA cluster *C*_*a*_, it would be excluded from the computation. Thus, the final distance between an HLA cluster and a disease-associated HLA allele is obtained as the weighted sum of pairwise distances between all HLA alleles in the cluster and the risk HLA allele.

### The definition of the THNet MS score

The THNet Matching Score (MS) quantifies the HLA distance between a donor and a recipient by evaluating the donor-specific HLA alleles, which are alleles that appear only in the donor but are absent in the recipient. If no donor-specific HLA alleles exist, the THNet MS is set to zero. For class I HLA alleles, HLA-A, HLA-B, and HLA-C are considered, while for class II HLA alleles, only HLA-DRB1 and HLA-DQB1 are included, as these have been shown to be most associated with transplantation outcomes in previous studies. The distance between the donor-specific HLA allele set D and the recipient HLA allele set R is calculated as:

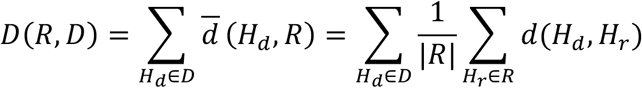

where *D*(*R*, *D*) represents the overall HLA distance between the donor *D* and the recipient *R*, *d*(*H*_*d*_, *R*) represents the mean distance of each donor-specific HLA allele *H*_*d*_ to all recipient HLA alleles, *d̅*(*H*_*d*_, *H*_*r*_) is the pairwise HLA distance between donor-specific allele *H*_*d*_ and recipient allele *H*_*r*_, |*R*| is the number of HLA alleles in the recipient. Thus, THNet MS is defined as the sum of the average distances of all donor-specific HLA alleles to recipient HLA alleles. For class I HLA alleles, mismatches are calculated across HLA-A, HLA-B, and HLA-C, while for class II HLA alleles, the pairwise distance is restricted within HLA-DRB1 and HLA-DQB1.

### Eplet score and combined score

The eplet scores of the organ transplantation data were calculated using the HLAMatchmaker service(*118*), specifically the ABC Eplet Matching Program V4.0 for class I HLA alleles and the DRDQDP Eplet Matching Program V3.1 for class II HLA alleles. The number of antibody-verified eplets was used as the eplet score for both class I and class II HLA loci. For HSCT transplantation, the eplet scores were obtained directly from their original study(104). To facilitate comparison, both the MS score and the eplet score were normalized to the same scale using z-score transformation. The combined score was then defined as the maximum of the two scores for each transplantation entry, based on the assumption that T-cell-mediated and B-cell-mediated graft rejection operate in a competitive relationship.

### Transplantation data

This study used data from the Scientific Registry of Transplant Recipients (SRTR)(*103*). The SRTR data system includes data on all donor, wait-listed candidates, and transplant recipients in the US, submitted by the members of the Organ Procurement and Transplantation Network (OPTN). The Health Resources and Services Administration (HRSA), U.S. Department of Health and Human Services provides oversight to the activities of the OPTN and SRTR contractors. In addition to the organ transplantation dataset, we also incorporated data from a hematopoietic stem cell transplantation (HSCT) dataset provided by the Center for International Blood and Marrow Transplant Research (CIBMTR)(*104*). The CIBMTR dataset is primarily funded by the Public Health Service (U24CA076518) from the National Cancer Institute, as well as grants from the National Heart, Lung, and Blood Institute; the National Institute of Allergy and Infectious Diseases; HRSA (75R60222C00011); the Office of Naval Research (N00014-23-1-2057, N00014-24-1-2507); the National Marrow Donor Program (NMDP); and the Medical College of Wisconsin. To ensure data consistency and relevance, we applied the following filtering criteria: (1) both the donor and recipient must have complete HLA information for HLA-A, HLA-B, HLA-C, HLA-DQB1, and HLA-DRB1; (2) samples must not contain any unpredictable HLA alleles; (2) the donor-recipient pair must have at least one mismatched HLA allele; and (4) recipient outcome data must be available. Additionally, for the SRTR dataset, four-digit HLA alleles were inferred based on the most frequent allele within each corresponding two-digit HLA superfamily in the White population(*119, 120*).

### SRTR data disclaimer

The data reported here have been supplied by the Hennepin Healthcare Research Institute (HHRI) as the contractor for the Scientific Registry of Transplant Recipients (SRTR). The interpretation and reporting of these data are the responsibility of the author(s) and in no way should be seen as an official policy of or interpretation by the SRTR or the U.S. Government.

### Statistical test

All significance assessments in the boxplots throughout the paper were performed using Welch’s t-test, which assumes normality but does not require equal variance between the two comparison groups. Additionally, all significance assessments in the scatterplots were calculated using Pearson correlation.

### Code availability

The THNet model is publicly available on GitHub (https://github.com/Mia-yao/THNet) and PyPI (https://www.piwheels.org/project/thnet/). It provides two main functionalities: HLA typing inference based on the TCR repertoire and MS score calculation. The 9 million TCR-HLA pairs exhibiting significant co-occurrence patterns has been deposited in Zenodo (10.5281/zenodo.14814254).

## Supplementary Materials

Fig. S1. Identification of HLA-associated TCR clusters.

Fig. S2. Most HLA models achieve a high F-score

Fig. S3. HLA pairs with less distances exhibit more similar peptide-binding capacities

Fig. S4. Class II HLA-based population clustering

Fig. S5. HLA cluster-disease association is not primarily driven by HLA haplotypes

Fig. S6. Influence of donor and recipient ethnic background on kidney transplant outcomes

Fig. S7. The THNet mismatch score is independent from the eplet score

Fig. S8. The class I THNet Mismatch Score (MS) is associated with the survival of organ transplant patients

Table S1: Predictable HLA allele list for class I HLA and class II HLA loci

Table S2: The distance between each class I HLA allele pair

Table S3: The distance between each class II HLA allele pair

Table S4: The disease-associated HLA clusters in White population Table S5: The disease-associated HLA clusters in Asian population Table S6: The disease-associated HLA clusters in Black population

