## Supplementary figures for "Quantitative and large-scale investigation of human TCR-HLA Cross-Reactivity"

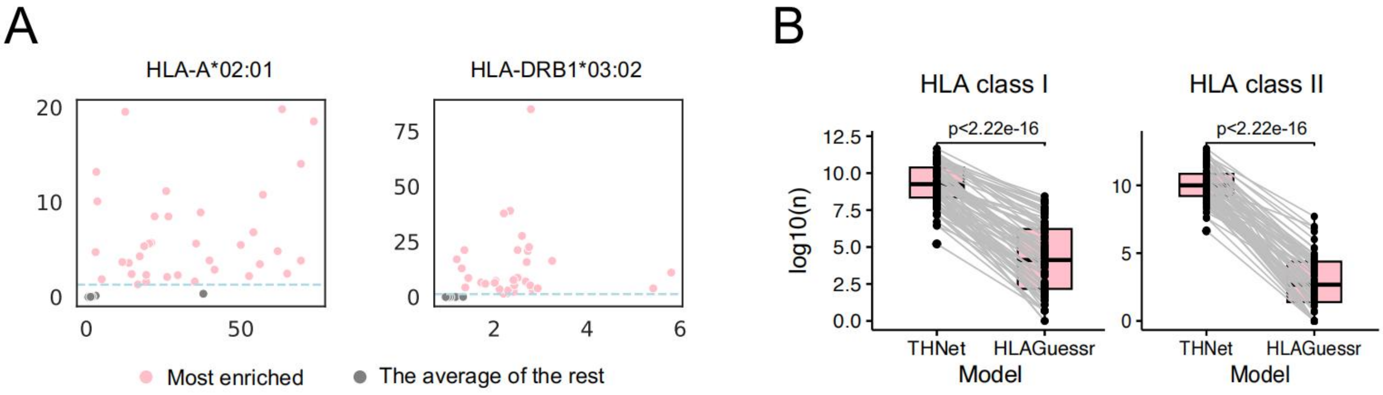


**Figure S1. Identification of HLA-associated TCR clusters.**

(A) The odds ratio and negative $\log_{10}$-transformed *p*-value of V genes from HA TCRs paired with different CDR3 regions. The pink dot indicates the most enriched V gene–CDR3 combination, while the grey dot represents the second most enriched combination. Examples are shown for HLA-A*02:01 (class I HLA) and HLA-DRB1*03:02 (class II HLA). The light blue dashed line represents the threshold where $p = 0.05$. (B) The $\log_{10}$-transformed number of HA TCRs used in THNet and HLAGuessr. Each black dot represents an HLA allele, with the same alleles connected by a grey line between the two models.

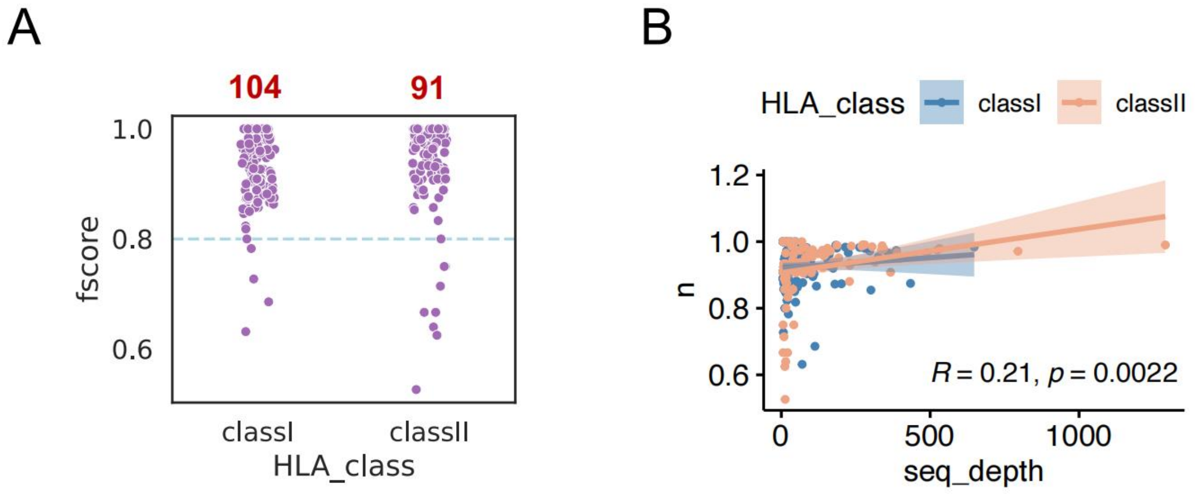


**Figure S2. Most HLA models achieve a high F-score**

(A) The F-score of each HLA allele calculated on the whole combined dataset. The light blue dashed line represents the threshold where F score= 0.8. Each dot represents an HLA allele model. (B) The association of F-score (y-axis) and the number of positive samples (x-axis) of each HLA allele model. Each dot in the plot represents an HLA allele model. The 95% confidence interval is indicated. Class I HLA and class II HLA were colored in blue and orange respectively.

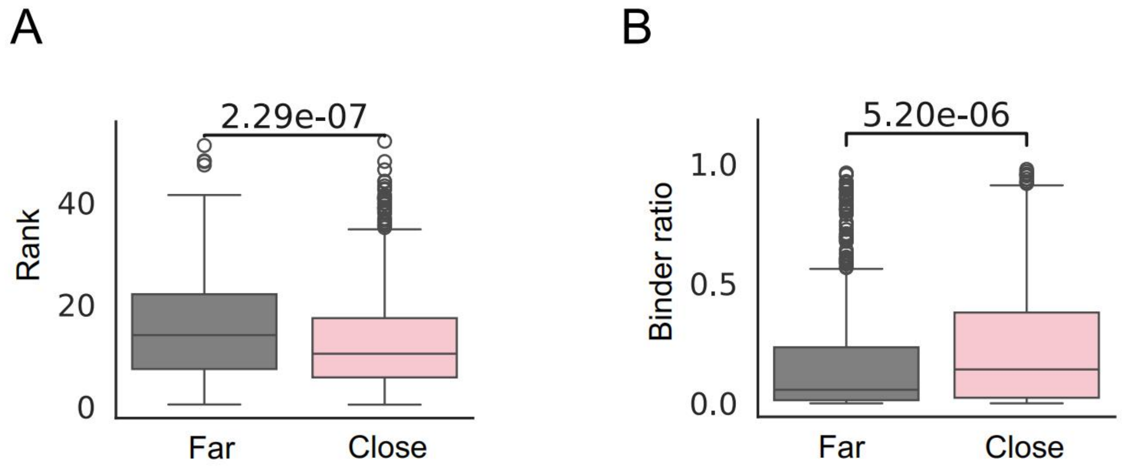


**Figure S3. HLA pairs with less distances exhibit more similar peptide-binding capacities**

(A) Peptide binding ranks of known HLA-binding peptides compared between their closely related and distantly related HLA counterparts. (B) The proportion of binding peptides among known HLA-binding peptides when compared between close and distant HLA counterparts. Close HLA pairs are defined as those with a distance in the bottom 10th percentile, while far HLA pairs are defined as those with a distance in the top 90th percentile.

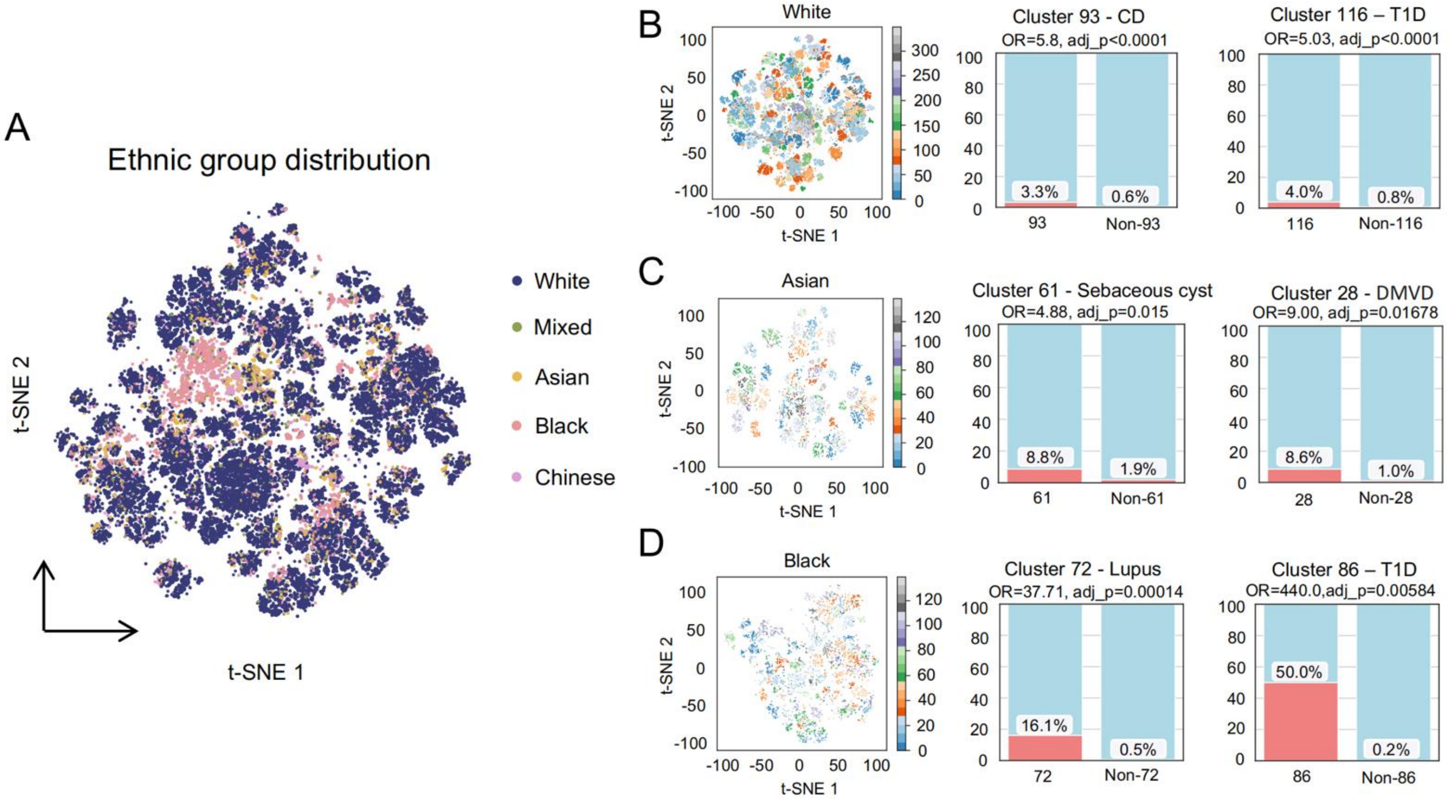


**Figure S4. Class II HLA-based population clustering**

(A) t-SNE plot depicting population clustering in the UK Biobank dataset based on class II HLA composition. Samples are colored according to race annotations in the UK Biobank database. Each dot represents one or more samples sharing the same HLA class II genotype. If a genotype is shared by samples from different racial backgrounds, the color is assigned based on the most frequent race within that genotype. (B–D) t-SNE plots displaying population clustering for White, Asian, and Black British populations, accompanied by stacked bar plots showing Fisher's exact test results for the two clusters with the most significant disease associations within each population. The x-axis of the bar plots indicates samples with/without the HLA cluster, while the y-axis represents the percentage of samples with specific diseases (CD: celiac disease; T1D: type 1 diabetes; DMVD: Degenerative mitral valve disease).

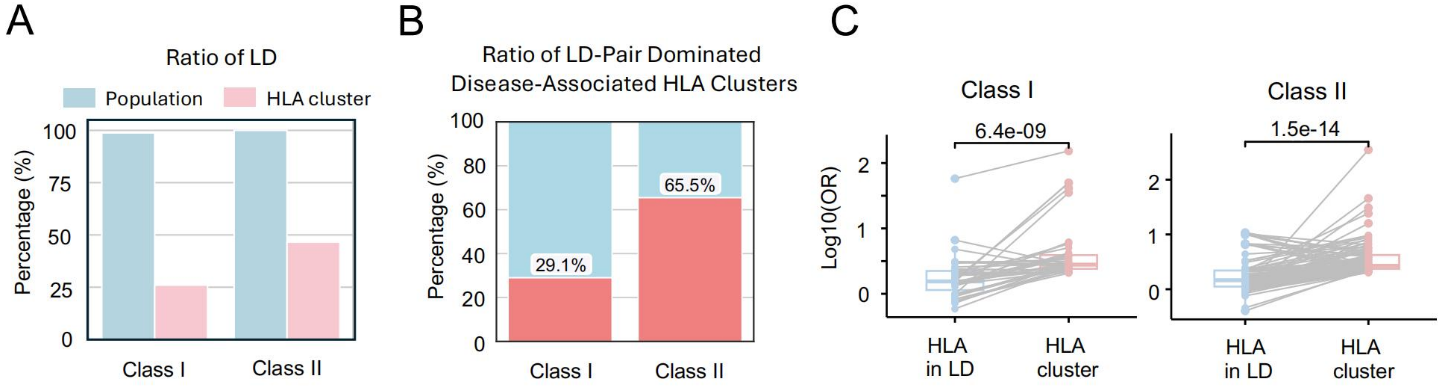


**Figure S5. HLA cluster-disease association is not primarily driven by HLA haplotypes**

(A) Bar plot showing the proportion of HLA allele pairs in linkage disequilibrium (LD) at the individual level and within HLA clusters for both class I and class II HLA alleles. The blue bars represent the percentage of individuals carrying at least one HLA pair in LD, while the pink bars indicate the proportion of HLA clusters dominated by HLA pair in LD. (B) Stacked bar plot depicting the proportion of significantly disease-associated HLA clusters dominated by HLA allele pairs in LD for class I and class II HLA alleles. (C) Box plot comparing the disease odds ratios (OR) of HLA LD-dominated disease-associated HLA clusters with their corresponding HLA pair in LD.

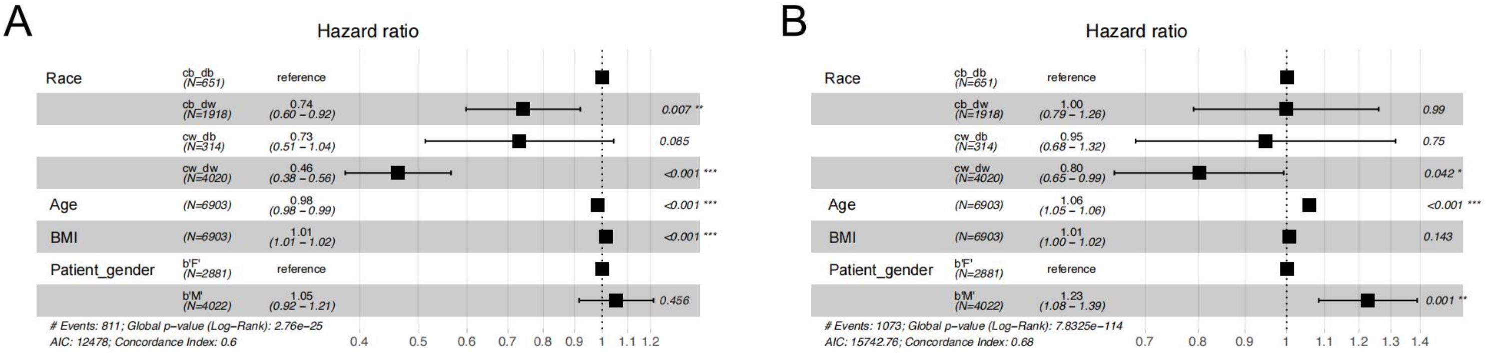


**Figure S6. Influence of donor and recipient ethnic background on kidney transplant outcomes**

(A) Cox proportional hazards model illustrating hazard ratios and *p*-values for kidney transplant rejection, analyzing the ethnic background of donors and recipients as risk factors. The model is adjusted for potential confounders, including age, body mass index (BMI), and gender. F: female; M: male. Categories are defined as follows: 'cb_db' indicates both donor and recipient are Black; 'cb_dw' indicates the recipient is Black and the donor is White; 'cw_db' indicates the recipient is White and the donor is Black; and 'cw_dw' indicates both donor and recipient are White. (B) Cox proportional hazards model illustrating hazard ratios and *p*-values for patient survival outcomes following kidney transplantation, with donor and recipient ethnic background analyzed as risk factors. The model is adjusted for potential confounders, including age, body mass index (BMI), and gender.

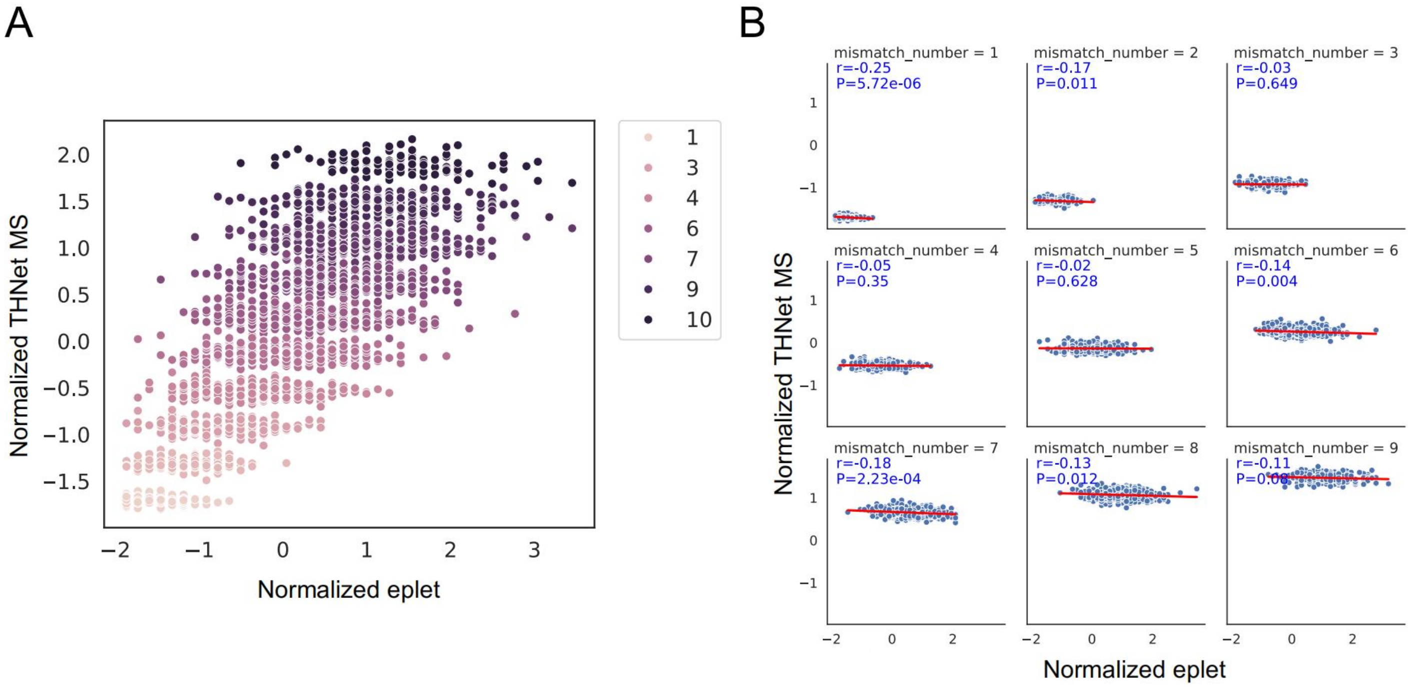


**Figure S7. The THNet mismatch score is independent from the eplet score**

(A) Scatterplot showing the correlation between normalized THNet mismatch score (y-axis) and normalized eplet score (x-axis), colored by the number of HLA mismatches. (B) Scatterplot showing the correlation between normalized THNet MS (y-axis) and normalized eplet score (x-axis), conditioned on mismatch numbers ranging from 1 to 9. Concordance and statistical significance were assessed using Pearson correlation.

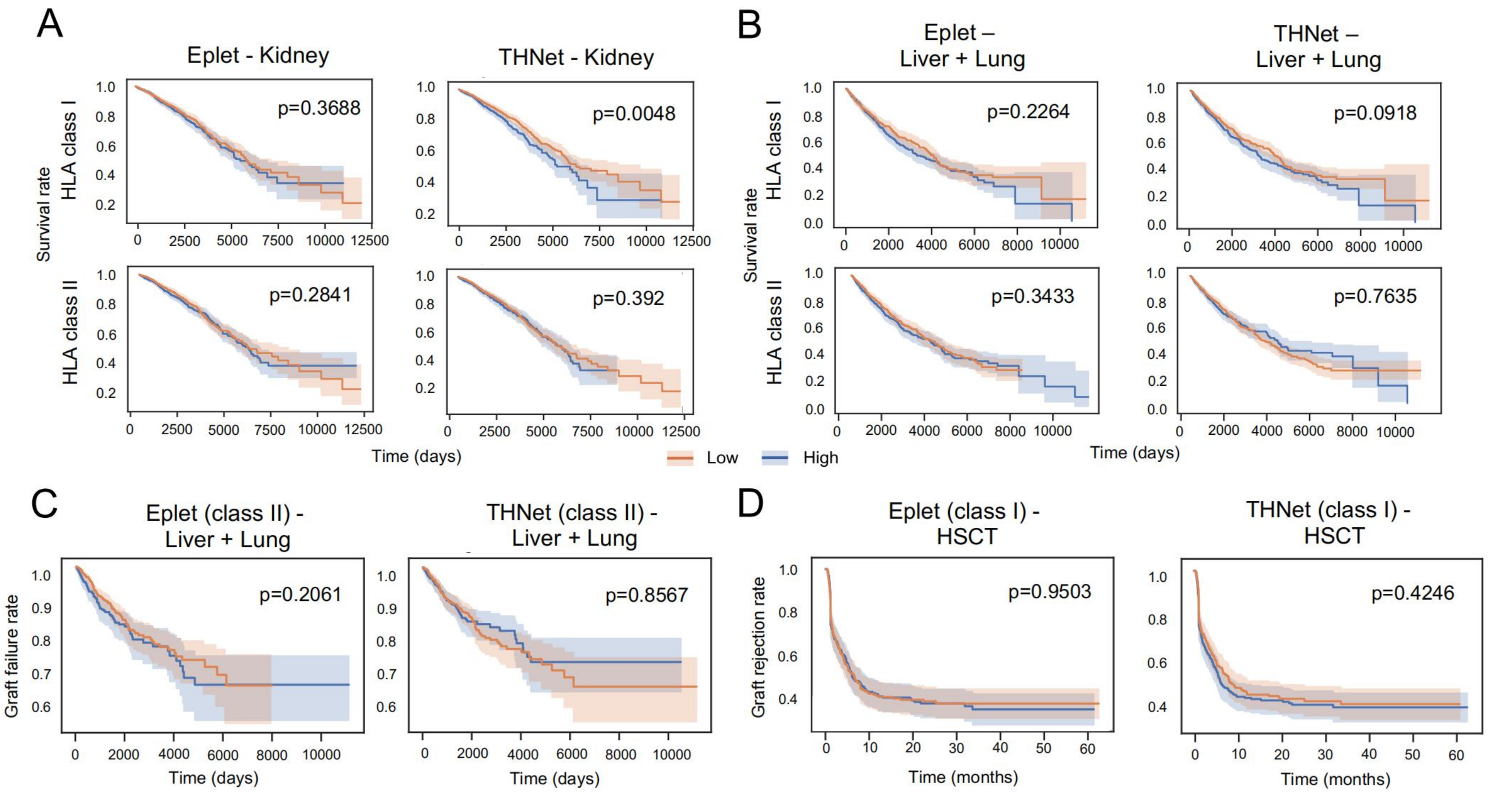


**Figure S8. The class I THNet Mismatch Score (MS) is associated with the survival of organ transplant patients**

(A) Kaplan-Meier curves depicting kidney transplant patient survival rates using the eplet score and THNet MS as indicators, with results for class I and class II HLA scores presented in two rows. Statistical significance was assessed using the log-rank test. (B) Kaplan-Meier curves depicting liver and lung transplant patient survival rates using the eplet score and THNet MS as indicators, with results for class I and class II HLA scores presented in two rows. (C) Kaplan-Meier curves showing liver and lung transplant failure rates for patients grouped by low/high class II eplet score and class II THNet MS. (D) Kaplan-Meier curve showing hematopoietic stem cell transplantation (HSCT) rejection rates for patients grouped by low/high class I eplet score and class I THNet MS.
